# Novel Three-Dimensional Organoid Sarcoidosis Granuloma Model

**DOI:** 10.1101/2020.09.05.284398

**Authors:** Tess M. Calcagno, Chongxu Zhang, Runxia Tian, Babak Ebrahimi, Mehdi Mirsaeidi

## Abstract

Sarcoidosis is a multi-system disorder of granulomatous inflammation which most commonly affects the lungs. Its etiology and pathogenesis are not well defined in part due to the lack of reliable modeling. This article presents the development of a novel *in vitro* three-dimensional lung-on-chip organoid designed to mimic granuloma formation. A lung on chip fluidic macrodevice with three channels for cell culture insertion was developed and added to the previously developed a lung-on-membrane model. Granulomas were cultured from blood samples of patients with sarcoidosis and then inserted in the air-lung-interface (ALI) of the microchip to create a three-dimensional organoid sarcoidosis model (OSGM). The model was tested for cell viability with fibroblasts. We measured the cytokine profiles in medium of OSGM and compared with lung model without granuloma. Concentration of IL-1beta, Il-6, GM-CSF, and IFN-gamma were found significantly higher in OSGM group. The current model represents the first 3D OSGM created by adding a microfluidics system to a dual-chambered lung on membrane model and introducing developed sarcoid-granuloma to its ALI.

## Introduction

### Sarcoidosis

Sarcoidosis is a heterogenous multisystem disorder characterized by the formation of noncaseating granulomas which can lead to extensive fibrosis. Its etiology is still largely unknown, though genetic, environmental, and infectious causes have all been considered [1]. Its clinical presentation is extremely variable ranging from an asymptomatic incidental finding to a life-altering disease. Although sarcoidosis can virtually affect any organ, pulmonary sarcoidosis is the most prevalent and is associated with the highest morbidity and mortality, usually related to pulmonary hypertension and extensive pulmonary fibrosis.

The features of human pulmonary sarcoidosis that could be modeled are still undefined, largely due to the limited understanding of etiology and pathogenesis of sarcoidosis. Human exposure to a wide variety of infectious and environmental components may cause different sarcoidosis phenotypes. The predisposition susceptibility to sarcoidosis is an important consideration in developing an acceptable animal model, which is currently not well developed. In addition, in some cases, it is difficult to translate experimental findings in animal models to humans. Inter-species variability and associated limitations of animal models have been widely established [2]. In addition, traditional cell line culture methods suffer from severe limitations [3]. Organ-on-a-chip technology combines tools from microfluidics and instrumentation control with advances in cell and tissue engineering to engineer biomimetic models of biological systems with high fidelity[4-7].

Two-dimensional *in vitro* cell culture models combined with *in vivo* animal testing served as gold standard methods for decades of medical advancements in pulmonary research [8]. However, results from experiments in animals cannot always be reliably translated to human disease processes due to interspecies differences in physiology and genetics, especially in the setting of inflammatory and host-pathogen response [9]. In two-dimensional (2D) cell culture, cells of one type can be studied in a low-cost standardized fashion which allows for massive high throughput screenings and experimental protocols which can easily be replicated. However, 2D cell culture is limited by its fixed device architecture and stagnant culture media. Simplistic representation fails to recreate lung-specific microenvironments which rely on the interplay between multiple cell types within a complex physiologic system [10].

Contemporary three-dimensional (3D) lung on chip models optimize clinical advancement by more accurately reconstructing normal lung physiology. Lung on chip models use microfluid-based cell culture on a micro-or nano-sized chip allowing for perfusion of targeted cells into the chip and reduction of reagent consumption as compared to conventional cell culture technique. Most lung-on-chip models (LOCM) are designed to mimic the environment of the alveolar air-blood barrier comprised of alveolar epithelial cells and vascular endothelial cells in the setting of applied mathematical models which consider both fluid mechanics and stretch frequency [11]. However, microfluid-based cell culture technology used in lung on chip modeling is limited by non-standardized culture protocols and low throughput capacities.

In the setting of sarcoidosis, novel 3D models properly mimicking granuloma formation have not yet been created; this has contributed to the lack of available targeted therapies. A novel lung on chip platform has been developed by adding a microfluidics system to the previously developed dual-chambered lung on membrane model. Developed granulomas were then introduced to the air-lung interface, making this the first three-dimensional lung on chip model mimicking sarcoidosis.

## Results

Mature granuloma developed within 3 days as shown in Figure 1. The macrophages and lymphocytes were aggregated and formed a large structured granuloma.

**Figure 1.**
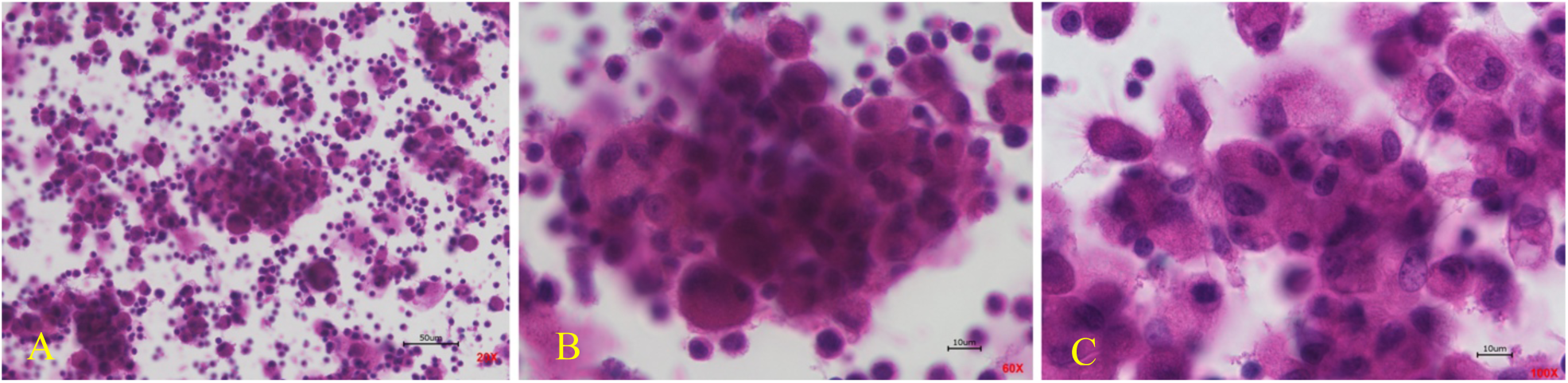
*In vitro* granulomas from PBMC taken from blood of patients with sarcoidosis. **A** displays granuloma at 20X magnification, **B** displays granuloma at 60X magnification, **C** displays granuloma at 100X magnification.

The fluidic macrodevice was developed and shown in Figure 2.

**Figure 2.**
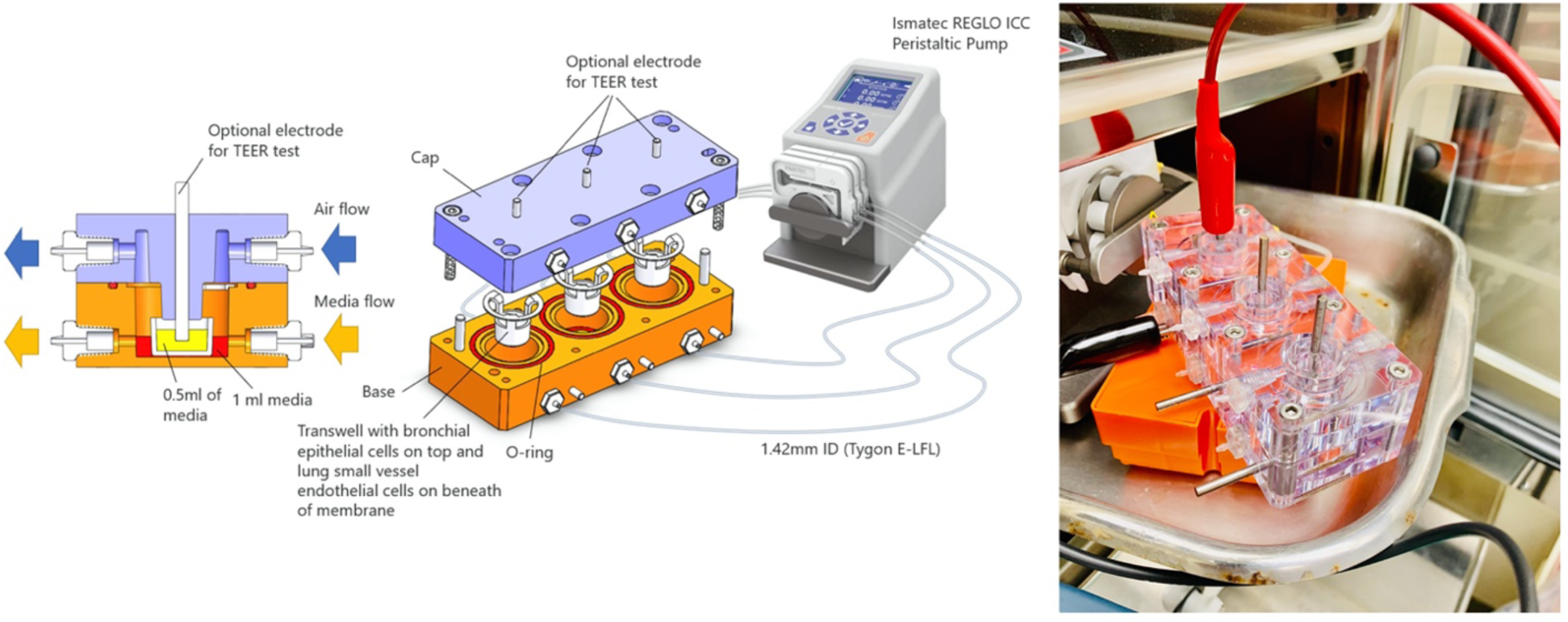
Design and fabrication of fluidic macrodevice and physical device in action

The lung on chip model was developed as shown in Figure 3 which displays TEM images of the lung model and its varying components (NHBE cells, endothelial cells, and intervening membrane).

**Figure 3.**
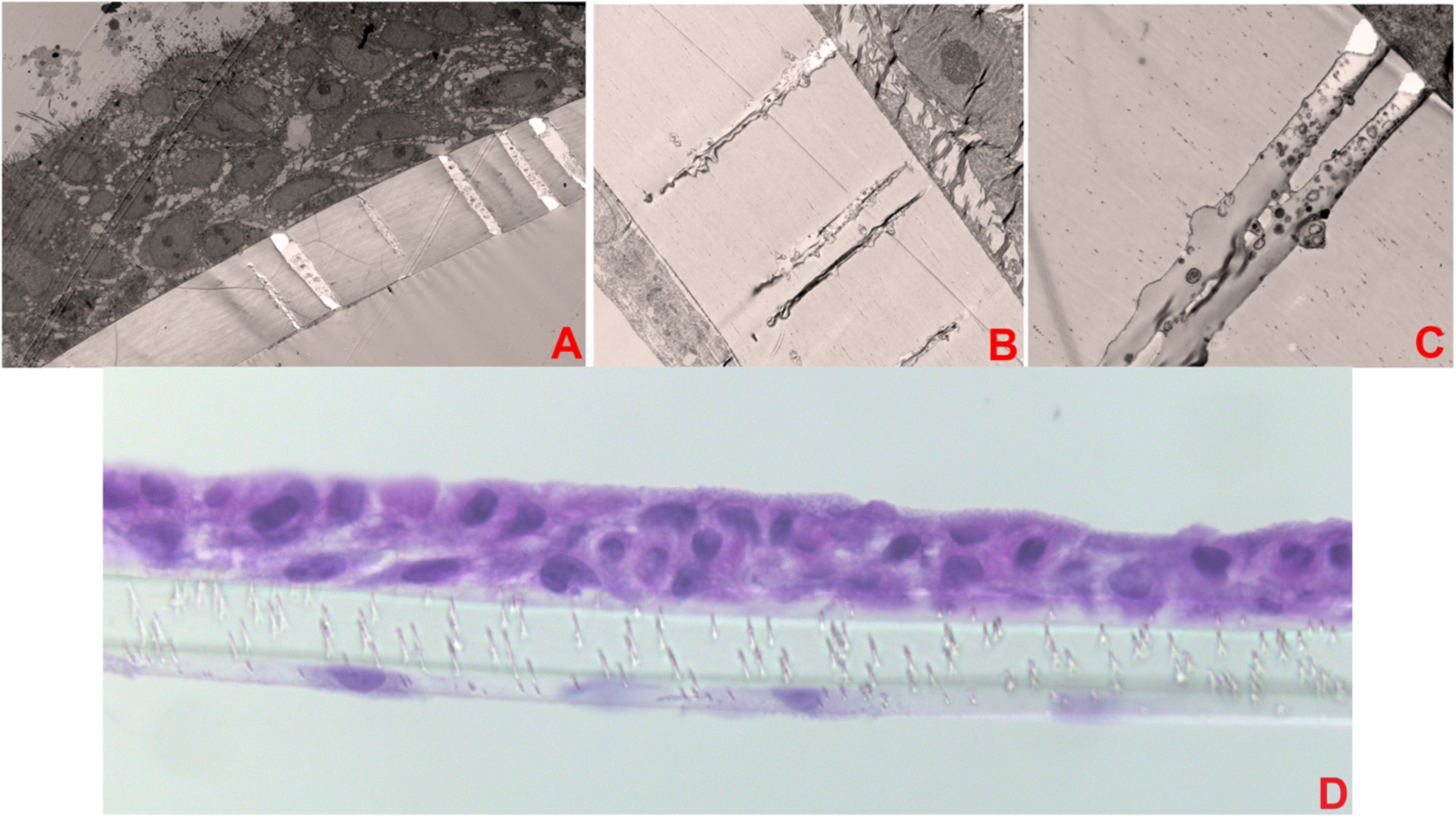
Displays images of dual-chambered lung on membrane model (LOMM). **A** shows LOMM with the red arrow pointing to NHBE cells, magnification X400, exposure 3000 (ms). **B** shows LOMM with the yellow arrow pointing to the polycarbonate membrane magnification X1500, exposure 3000 (ms). **C** shows LOMM with the purple arrow pointing to membrane with *0*.*4* μm pore magnification X4000, Exposure 3000 (ms). **D** shows LOMM displaying NHBE cells (red arrow), membrane (yellow arrow), and endothelial cells (green arrow), magnification X40.

Fibroblasts were cultured and immunofluorescence staining was done to test the functionality of the device as shown in Figure 4.

**Figure 4.**
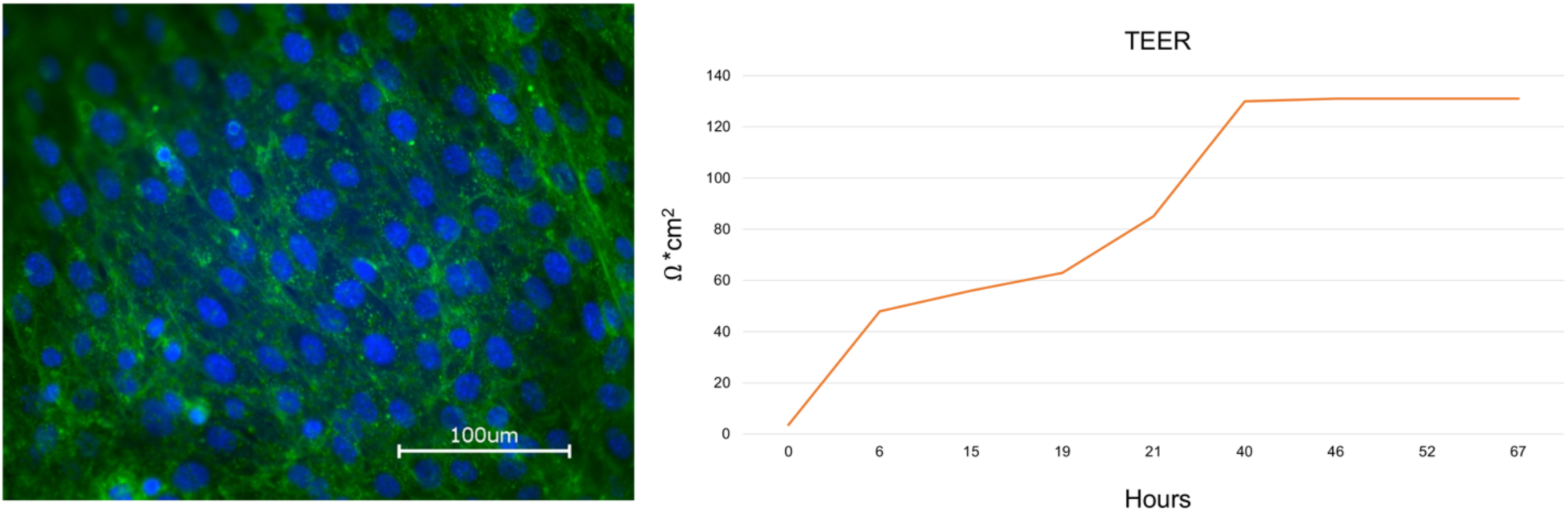
Shows immunofluorescence staining of fibroblasts on membrane inside the Lung on chip device with WGA staining. Right image shows TEER results that shows flattening after 40 hours.

The OSGM was formed; developed granulomas were introduced to the air-lung interface of the lung on chip model as seen in TEM images in Figure 5.

**Figure 5.**
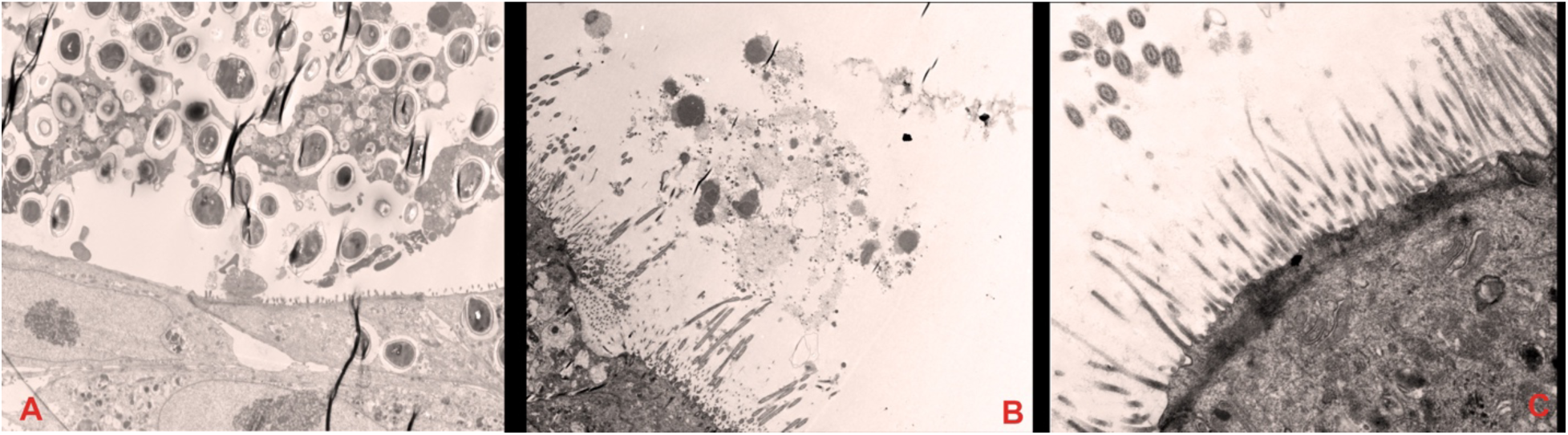
Shows transmission electron microscopic images of OSGM. **A** shows bronchial epithelial cell with cilia (Red arrow) and granuloma (lymphocytes and macrophages) with Blue arrow, magnification of X1200. **B** shows macrophages and lymphocytes from developed granuloma in the ALI, magnification of x1200, Exposure 3000 (ms). **C** shows mature bronchial epithelial cell with cilia (Blue arrow) and microvillia (Red arrow), magnification X5000. Red arrows show NHBE cells.

Cytokine measurements were taken in the OSGM as compared to control; there were statistically significant differences in IL-beta expression (P=0.001953), Il-6 expression (P=0.001953), GM-CSF expression (P=0.001953), and INF-gamma expression (P=0.09375) as shown in Figure 6.

**Figure 6.**
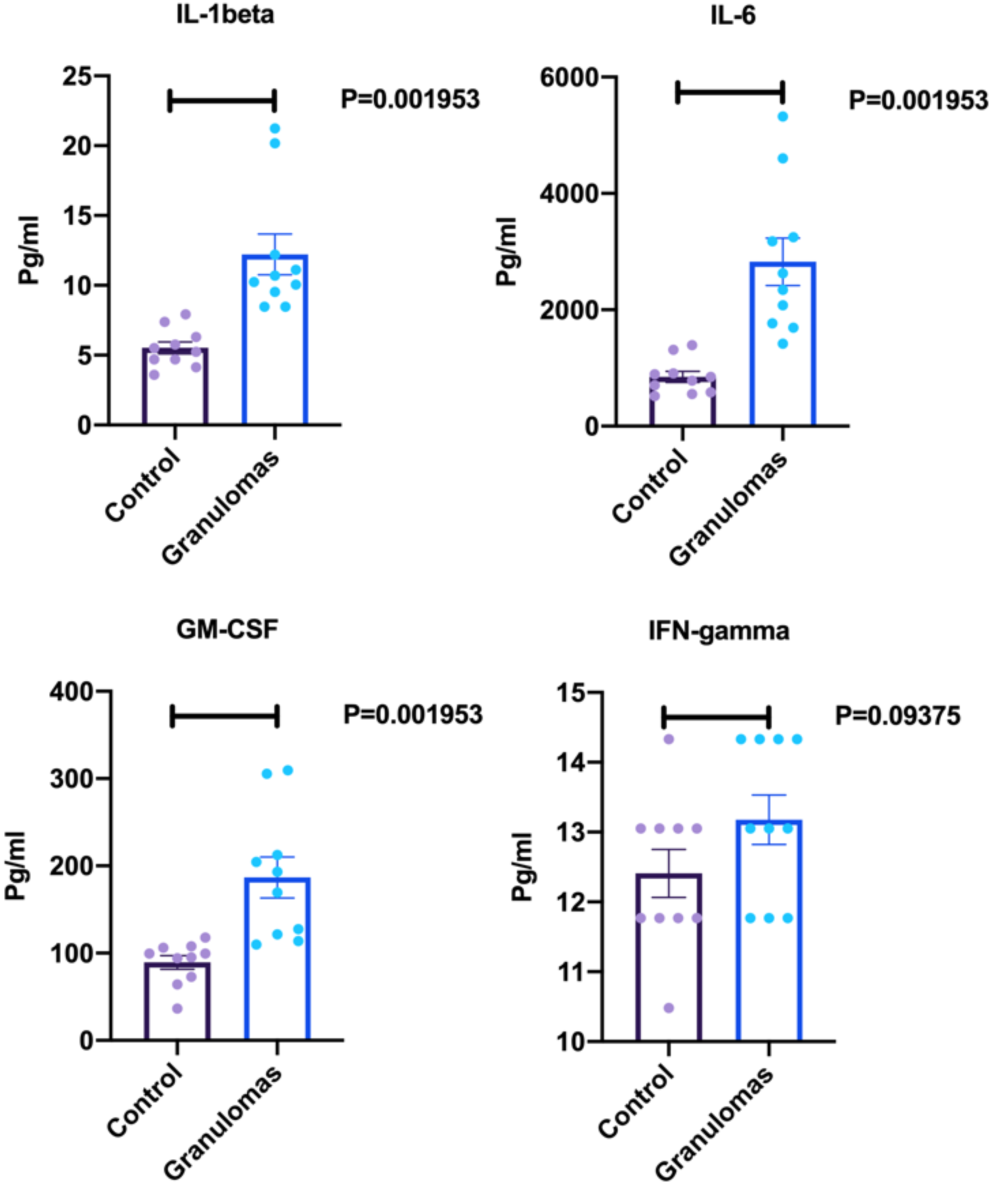
Displays ELISA results for cytokine response measure in pg/ml (IL-1beta, Il-6, GM-CSF, and IFN-gamma). Control represents the lung on chip model without granuloma in ALI. Granulomas represent lung on chip model with granulomas in ALI (n=10 for each group).

## Discussion

The first Three-Dimensional Organoid Sarcoidosis Granuloma Model (3D OSGM) is developed by adding a microfluidics system to our previously developed lung on membrane model and fully developed granulomas was introduced to its air-lung interface. The newly developed dual chamber lung on membrane model uses standardization techniques, and compatible to use artificial intelligence (AI) to combine the benefits of reproducibility from 2D cell culture with the complexity of lung on chip microfluid models. It is comprised of three individually controlled channels which are independently sealed and contain individual peristaltic pumps to control the flow rate of media in each channel with precision. The researcher can control the flow rate in real time using an iOS and Android compatible microcontroller. Microfluid design is used with 1 mL of media in each chamber; and cell cultures are inserted via a 12mm Transwell cup with a 0.4 μm pore Polyethylene Terephthalate membrane insert. Further standardization is permitted with insertion of six electrodes which monitor continuous electrical impedance measurement (TEER test) every 60 minutes. Our model is also efficient; it is capable of performing three different experiments at the same time with up to four different cell lines.

Lung on chip (LOC) models emulate lung physiology using a microfluid design combined with culture inoculation [12]. Huh et al. developed the first LOC model which contains two microchannels coated with endothelial cells and alveolar epithelial cells separated by a 10um collagen coated polydimethylsiloxane (PDMS) porous membrane. Two separate channels with vacuum applied suction surround the lateral aspects of the model to stimulate negative intrapleural pressure involved in inspiration [4]. Humayun et. al added a hydrogel micro layer which allowed for the development of a LOC model with smooth muscle and epithelial cells to mimic bronchiolar microenvironments [13]. LOC models have since been used successfully to mimic lung injury, lung inflammation, pulmonary fibrosis, and lung cancer, but not sarcoidosis [14-17].

Prior to the development of our model, there were no 3D lung models designed to mimic sarcoid-like granulomatous disease. The etiology of sarcoidosis is poorly understood, but it is likely a gene-environment interaction which triggers granuloma formation via antigen presentation, CD4 ^+^ T Cell activation, and secretion of cytokines including interferon-γ, tumor necrosis factor (TNF)-α, and transforming growth factor-β [18]. The inability to recreate sarcoid granulomas *in vivo*, has severely limited our ability to discover novel pharmacologic therapies, characterize its etiology, and understand its heterogenous clinical presentation.

In previous investigations, sarcoid has been studied using biopsies, animal models, and single cell lines. Biopsies of granulomas from diseased patients have been used to characterize sarcoid histologically at a stagnant point in time, but they cannot properly represent the dynamic interplay between multiple cell types seen in sarcoid-like granulomas [19]. Many animal models made to simulate granuloma formation have been developed in the past ten years, but development is limited since sarcoidosis does not occur naturally in most animals. In order to simulate sarcoid-like granulomas in animals, animals are exposed to environmental antigens [20]. For example, Werner et al. created a pulmonary granulomas in mice by introducing *Propionibacterium acnes*, an environmental bacterium possibly implicated in the development of sarcoidosis in human lung tissue [21]. Such models have been limited by lack of similarity to polygenic human sarcoid-induced granulomatous processes involving the interplay between T-cell, cytokine, and aberrant macrophage-mediated immune responses [22]. Two-dimensional cell cultures using BAL or blood samples have been used to study antigenic response in a single cell population, however this fails to account for a sarcoid microenvironment incorporating multiple cell types [19].

The current model is designed based on mycobacteria component which is one of the potential etiologies of sarcoidosis. However, it does not consider granuloma formation in response to other potential environmental exposures. This may limit future study to one pathophysiologic subset of sarcoid disease.

### Future Directions

Our new model which uses three-dimensional structuring, compatible to AI, and standardization techniques is the first of its kind to simulate sarcoidosis. AI will be the next step of development of this device. AI can control TEER testing, volume control as well as adding new medication of chemical to the experimental chamber. This development circumvents a large hurdle in the progression of sarcoidosis research. The model is scalable to be incorporated with four cell groups including fibroblast and circulatory macrophages. It will allow for better characterization of a heterogenous and complex multi-organ disease. Furthermore, efficient testing of novel pharmacotherapeutic agents will bring agents into clinical trials more quickly.

## Methods

### Development of Granuloma

Peripheral blood mononuclear cells (PMBCs) were isolated from peripheral blood samples of patients with confirmed sarcoidosis taken from University of Miami Sarcoidosis biobank as previously described [23]. A quantity of 2 x10^6^ PBMC was cultured in a 12-well tissue culture dish and challenged with in-house microparticles developed to induce granuloma formation (Figure 1) [24].

### Design and fabrication of fluidic macrodevice

The lung on chip fluidic macrodevice is comprised of two main components, base and top cover (cap). Both components are CNC machined Polycarbonate blocks that sealed together to create a multi-channel platform with independent flow-control capability. There are three channels at the base, where cell cultures are inserted via membrane dual cell culture plate (12 mm Transwell with 0.4 μm pore polycarbonate membrane insert) (Figure 2). Channels are independently sealed via Soft Viton® Fluoroelastomer O-Ring (Made of FDA Listed Material, SAE AS568). For each channel at the base, there is a corresponding channel in the top cover, providing controlled air supply to the cell culture. Ismatec REGLO ICC peristaltic pump (3 channel/3 roller) was utilized to independently control the flow rate of the 3 channels for blood flow and single channel for air flow. If needed, a separate pump could be used to achieve independent air flow control. Masterflex Tygon E-LFL tubes with 1.42mm ID were used for fluid transfer. Considering 1ml blood volume in each chamber and 700mm tube length per channel, the pump speed was set to 20 rpm to achieve 2ml/min flow rate per channel resembling lung cell exposure to blood based on a cardiac output equal 5-6 liter/min. 6 electrodes (3 in the top cover and 3 in the base) were used for continuous electrical impedance measurement (TEER test).

### Development of lung on chip

We previously developed a lung-on-membrane model (LOMM) with a dual chamber including normal bronchial epithelial (NHBE) cells and human microvascular endothelial cells (Figure 3) [25]. We added a microfluidics system to the LOMM to develop a novel lung on chip.

To demonstrate functionality of the device, mice pulmonary fibroblasts were cultured due to their high rate of replication. Fibroblasts were cultured as previously described. As a summary, newborn mouse lung fibroblasts (Mlg 2908) were obtained from the American Type Culture Collection, Manassas, VA and cultured in Eagle’s minimal essential medium. TEER test was performed using the device‘s electrodes. The final TEER values were normalized but subtracting the baseline measurement from the read for each time spot and multiplied in the area of cell culture (1.12 cm^2^ for 12 well Transwell). The final value was reported as ohm/cm^2^.

Immunofluorescence (IF) staining was performed by using Wheat Germ Agglutinin (WGA). WGA was bought from Thermo Fisher Scientific (Catalog number: W11261), and manufacturer’s instructions were followed. Briefly, fixed cells were washed in PBS buffer 3 times; sufficient amounts of 5.0 μg/mL WGA labeling solution in HBSS buffer were then added to cover cells. The cells were then incubated for 15 minutes at room temperature. When labeling was complete, the labeling solution was removed, and cells were washed three times in HBSS buffer. Mounting Medium with DAPI (Vector lab, Catalog number: H-1200-10) was applied before taking image.

### Adding granuloma to air-lung interface (ALI)

A narrow scratch was made on the ALI and developed granulomas from 2 x10^6^ PBMC in 50 μL were directly added to ALI of lung on chip microchip to develop three-Dimensional Organoid Sarcoidosis Granuloma Model (3D OSGM).

### Cytokine measurements

To demonstrate the functionality of the device, experimental study was conducted. Ten LOMM and ten 3D OSGM were developed as previously described. The circulatory media were collected after 48 hours and ELISA testing was performed to measure cytokine response difference between LOMM with 3D OSGM. Elisa was performed using a kit from Invitrogen (Carlsbad, CA, USA) (Procartaplex human th1/th2 cytokine panel 11 plex from Invitrogen, cat # epx110-10810-901) per the manufacturers’ instruction.

## Author Contributions

Conceptualization: M.M.; Device design and development: M.M., B.E.; ELISA test: C.Z, R.T.M.M. Writing – Original Draft Preparation: T.C.; Writing – Review & Editing, T.C., C.Z, R.T., B.E., and M.M.

## Conflicts of Interest

Dr. Babak Ebrahimi is Director of Engineering at Genix-Engineering. The other authors declare no conflict of interest.

## Notes

### Competing Interest Statement

The authors have declared no competing interest.

